# An AI-Powered Trisomy 21 Research Assistant

**DOI:** 10.64898/2026.06.08.730893

**Authors:** Sutanu Nandi, Zenitha Sundararajan, Marc Subirana-Granés, Joaquin M. Espinosa, Milton Pividori, Kelly D. Sullivan, Matthew D. Galbraith, James C Costello

## Abstract

Down syndrome, caused by trisomy 21, increases the risk of diverse co-occurring conditions. With more than 34,000 related publications indexed in PubMed as of early 2026, keeping pace with this expanding literature is challenging. While general-purpose large language models are widely used for information retrieval, they often rely on broad training data rather than specific evidence. Retrieval-augmented generation (RAG) improves rigor and reliability of responses by linking model outputs to source texts. In research, source texts are peer-reviewed articles. Standard implementations treat all manuscript sections equally, allowing background text to rank as highly as experimental results. To focus model outputs on experimentally supported responses, we developed the T21 Research Assistant, a section-aware RAG system that prioritizes Results sections to ground responses in primary experimental evidence. The system draws exclusively from 1,789 open-access Down syndrome publications from PubMed Central, including 327 NIH INCLUDE-funded studies, and uses a multistage pipeline for query validation, retrieval, reranking, synthesis, and citation verification. Built on NVIDIA Nemotron models, it generates structured, cited responses. Evaluation using expert-curated questions demonstrated strong performance, achieving a BERTScore F1 of 0.712 and recall of 0.758, comparable to or exceeding leading proprietary and open-source models. T21 Research Assistant is available at: https://bioinformatics.cuanschutz.edu/t21-res-assi/

## 1. Introduction

Down syndrome (Trisomy 21; T21) affects nearly all systems in the human body and elevates risk for an array of co-occurring conditions (e.g., congenital heart defects, immune dysfunction, thyroid disorders, leukaemia, autoimmune conditions, and early-onset Alzheimer disease, among many others) (Antonarakis et al. 2020; Santoro et al. 2021). At the same time, individuals with DS display a lower incidence of many solid tumours, hypertension, and allergic sensitization (Antonarakis et al. 2020). The research community uses both human clinical studies and experimental models, including mouse strains, iPSCs, and other cell lines, to investigate these conditions (Antonarakis et al. 2020; Russo et al. 2024). Studying the link between T21 and co-occurring conditions requires an interdisciplinary integration of a vast body of literature (Bull et al. 2022; Weijerman and de Winter 2010; Lott and Head 2019; Dierssen 2012; Mégarbané et al. 2009).

Among the many active research fronts in this rapidly expanding field, significant contributions have advanced understanding of the molecular, immune, and metabolic changes observed in DS, leading to new therapeutic opportunities in DS. Foundational genomic analyses established that T21 consistently activates the interferon transcriptional response across multiple cell types (Sullivan et al. 2016), and subsequent metabolomics work demonstrated that overdosage of chromosome-21-encoded interferon receptors dysregulates the kynurenine pathway, elevating immunosuppressive and neurotoxic tryptophan catabolites in plasma and cerebrospinal fluid (Powers et al. 2019); building on this, multi-omics integration has uncovered coordinated immunometabolic dysregulation that stratifies individuals with DS into clinically distinct subgroups (Gillenwater et al. 2024), and recent multi-omic analysis of plasma from over 400 people identified perturbed bile acid metabolism and hepatic dysfunction as additional systemic consequences of T21(Dunn et al. 2026). Translating these molecular insights to clinical practice, a Phase II trial of the JAK1/3 inhibitor tofacitinib demonstrated normalization of autoimmune biomarkers and clinical benefit across dermatological conditions in DS, establishing JAK/STAT signaling as a tractable pharmacological target (Rachubinski et al. 2024). Together, these contributions exemplify the breadth of ongoing T21 research and the challenge of tracking advances across this multidisciplinary literature.

With over 34,000 PubMed-indexed publications on Down syndrome as of early 2026, it is a challenge to stay updated on research findings across this vast body of literature. In recent years, large language generative AI models (LLMs) have been widely adopted to answer queries across many domains. Generative AI models are trained on vast general-purpose text and tend to generate responses biased toward commonly used words and phrases reflective of the broad and generalized model training, rather than drawing on the specific evidence in peer-reviewed, Down syndrome research. A further concern is that LLMs are known to generate plausible but non-existent references; a recent audit of 2.5 million biomedical papers identified 4,046 fabricated citations across 2,810 publications, with the rate of fabrication increasing 12-fold between 2023 and 2026 (Topaz et al. 2026). Providing the model with relevant context directly from the Down syndrome literature helps to focus answers on what can be experimentally supported, with every claim traceable to a public and peer-reviewed publication.

More broadly, agentic AI systems and large language models have been successfully deployed across diverse biomedical domains, including general-purpose multi-task biomedical reasoning (Huang et al. 2025), autonomous analysis of single-cell RNA sequencing data (Alber et al. 2026), agentic design of gene-editing experiments (Qu et al. 2025), self-verified gene-set analysis using curated biological databases (Wang et al. 2025), LLM-derived embeddings for single-cell biology (Chen and Zou 2024), automated conversion of research papers into interactive AI agents (Miao et al. 2025), and end-to-end autonomous AI research pipelines (Lu et al. 2026). These successes demonstrate that grounding AI systems in domain-specific data and structured agentic workflows substantially improves their reliability and utility for specialized scientific tasks.

Retrieval-augmented generation (RAG) connects a generative large language model to a curated database of domain-specific knowledge from research publications as a context at the time each question is asked (Lewis et al. 2020). Rather than relying solely on what the model learned during training, the system first retrieves the most relevant context from the database and then asks the model to generate its answer using only those chunks as evidence. Thus, every claim in the response can be traced back to a specific paper and section, making the answer verifiable. RAG systems have been applied across several biomedical domains with measurable improvements in answer accuracy over standard language model baselines (Amugongo et al. 2025; Liu et al. 2025), but their deployment for the specialised condition of Down syndrome, where the literature is multidisciplinary and spans both human and model organism research, has not been addressed.

Furthermore, RAG models that rely on simple text retrieval treat all sections of a research publication with the same weight. However, the structure of a manuscript is designed to convey different kinds of information with the Introduction section being used to provide background and set up the research problem, the Methods section describes the experimental design and materials in detail, the Results section reports the observed and measurable findings objectively, and the Discussion section provides for interpretation, limitations, and broader implications. A sentence from a paper’s Introduction that provides background context, such as *"Down syndrome is associated with increased Alzheimer disease risk"*, can score just as highly in a standard similarity search as a sentence from the Results section reporting the actual measurements behind that association, even though only the second contains direct experimental evidence. Prioritizing information from the Results, Discussion, and Methods sections will shape the answers generated from RAG models to answer questions with well supported results, rather than generalized knowledge statements, while simultaneously facilitating the precise validation and formal attribution of experimental observations.

Here, we present the T21 Research Assistant, a multi-agent RAG that retrieves information from a database of 1,789 open-access Down syndrome publications from PubMed Central, including 327 articles funded through the NIH INCLUDE (INvestigation of Co-occurring conditions across the Lifespan to Understand Down syndromE) Project, ranks them by section priority, and generates structured responses with inline citations through a multilevel agentic pipeline covering retrieval, ranking, writing, citation checking, safety monitoring, LLM-as-Judge quality assessment, and quality review. We evaluated the T21 Research Assistant against a 12-question benchmark written by experts in Down syndrome research, using semantic similarity, word overlap recall, and an independent AI judge to measure how well the system’s answers match the expert reference answers. The section-priority design and the evaluation framework introduced here are directly applicable to RAG system development in other specialised biomedical domains.

## 2. Methods

### 2.1. System Architecture

The T21 Research Assistant is built using LangChain (v0.3+) (“LangChain: Observe, Evaluate, and Deploy Reliable AI Agents” 2022) for language model integration and LangGraph for deterministic pipeline orchestration (**Figure 1**). The system comprises three integrated layers: (i) a data ingestion pipeline that retrieves full-text publications from PubMed Central and enriches them with metadata from the OpenAlex, NCBI efetch, and CrossRef APIs, segments full texts at section boundaries, and encodes them as dense vector representations; (ii) a RESTful web API that receives queries, executes the eight-stage, multi-agent RAG pipeline, and returns structured answers with inline citations; and (iii) a browser-based Streamlit interface for submitting natural language questions and reading annotated responses.

**Figure 1.**
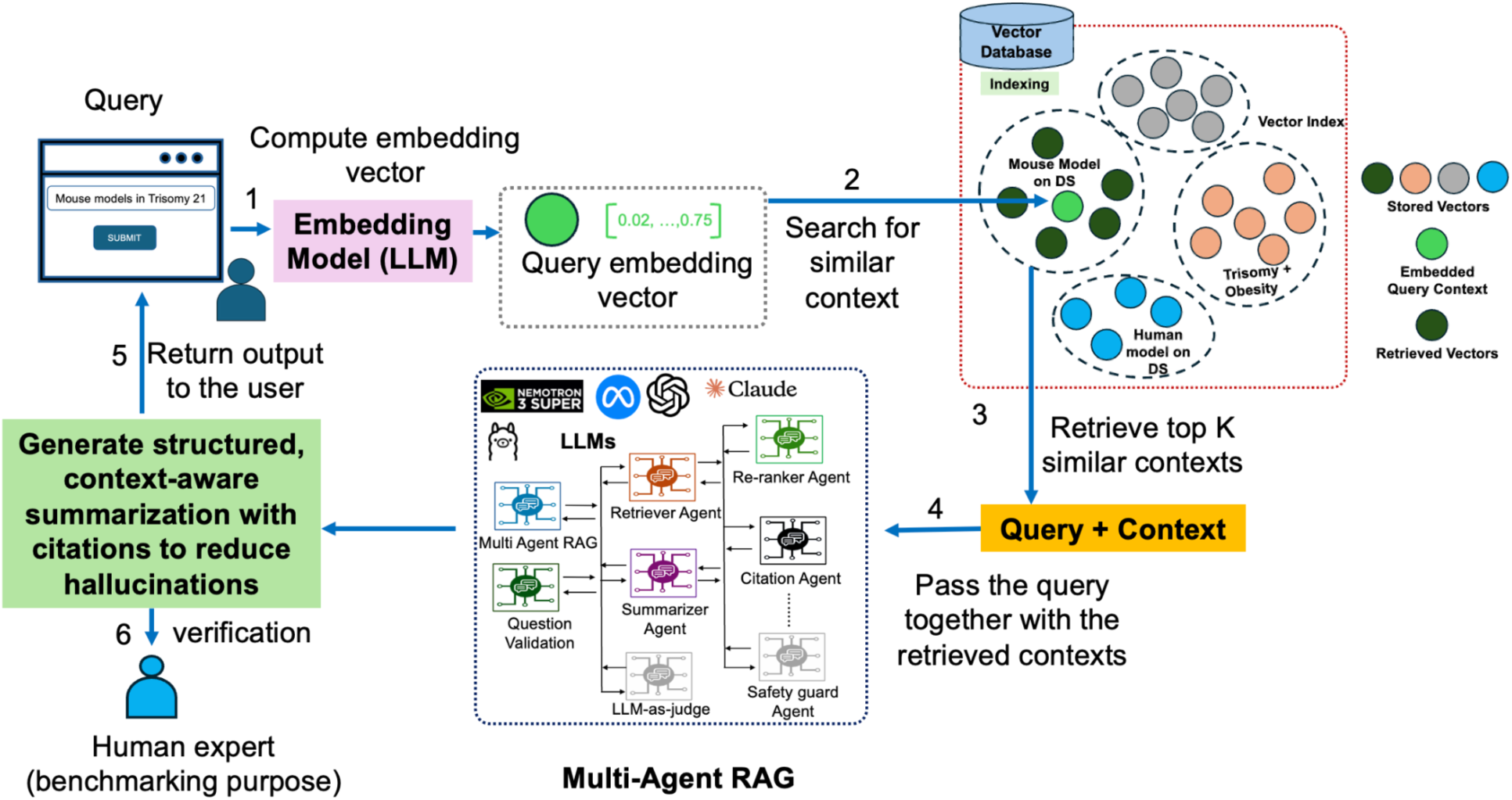
Overview of the T21 Research Assistant multi-agent RAG workflow. Steps are numbered sequentially. (1) A researcher submits a natural language query through the browser interface. (2) The query is converted to a 4,096-dimensional vector by the open source Nemotron embedding model (nvidia/llama-embed-nemotron-8b). (3) The query vector searches the Milvus vector database of 65,609 indexed INCLUDE publication chunks (IVF_FLAT index; nprobe = 64). (4) The top *k* most similar chunks are retrieved. (5) The original query and retrieved context pass through the eight-stage multi-agent LangGraph pipeline: Question Validation → Retrieve → Rerank (open source Nemotron cross-encoder) → Synthesize (open source Nemotron-3-Super; C-C-C prompt; temperature = 0.0) → Raw Evidence Extraction → Safety Guard → Finalize (citation hyperlinking; 150-word cap). LLM-as-Judge (automated quality scoring and guided rewrite). The pipeline generates a structured, context-aware summary with inline citations [PMCxxxxxxx:Section] and returns the output to the user. (6) Human expert evaluation against reference answers is used for benchmarking. LLM logos within the multi-agent box denote the supported chat providers (NVIDIA Nemotron, Meta Llama, OpenAI GPT, and Anthropic Claude).

A defining architectural choice is that the core computational stack relies entirely on open-source NVIDIA Nemotron models (NVIDIA Corporation 2026). User input queries are converted to 4,096-dimensional vector representations using *nvidia/llama-embed-nemotron-8b(Face 2025)* , an open source large embedding model served locally. Reranking is performed by *nvidia/llama-nemotron-rerank-1b-v2 (Face 2026)*, an open-source cross-encoder that scores candidate chunks for topic specificity and quantitative evidence. The default chat provider is Nemotron-3-Super, deployed via a remote Ollama server. Critically, the same *nvidia/llama-embed-nemotron-8b* model is used for all evaluation metrics (BERTScore and Semantic Similarity) and for UMAP visualisation, so the entire retrieval, synthesis, and evaluation pipeline operates within a single unified open source embedding space. This all-Nemotron default stack means that the system can be operated with no dependency on proprietary commercial API subscriptions. However, OpenAI (GPT-5.4), Anthropic (Claude Sonnet 4.6), and Google (Gemini) are supported as alternative or supplementary providers through a common configuration interface requiring no code changes.

### 2.2. Document Database and Processing

The corpus of text that supports the T21 Research Assistant comprises 1,789 open-access peer-reviewed publications on Down syndrome retrieved from PubMed Central (PMC). Articles were retrieved using the following PMC full-text search query: “"Down syndrome"[MeSH Terms] AND (open_access[Filter]) AND (published_article[Filter]) AND (pmc_public[Filter])”, with four filters applied: (i) Publication stage: Published journal article; (ii) Publication type: Case Reports excluded (PT ≠ Case Report); (iii) Collections: PMC Open Access Subset; (iv) Embargoed articles: Excluded . For each retained article, structured metadata were collected from four complementary APIs: (i) the OpenAlex API (Priem et al. 2022) provided bibliographic metadata including title, publication year, journal, language, authors, institutional affiliations, funders, and topic classification; (ii) the NCBI efetch API provided MeSH (Medical Subject Headings) controlled vocabulary terms, author keywords, and grant details(“NCBI Entrez Programming Utilities (E-Utilities)” 2009); and (iii) the PMC OAI-PMH interface (“The PubMed Central OAI-PMH API” 2001) provided structured full-text XML, parsed into six canonical sections: Abstract, Introduction, Methods, Results, Discussion, and Conclusion. Among these articles, 327 were funded through the NIH INCLUDE program and are identified by cross-referencing with the NIH INCLUDE Publications list (2017–2025); these articles are tagged with an INCLUDE provenance badge in the user interface. Together, the corpus spans publication years 1962–May 2026. Articles were segmented at section boundaries (Abstract, Introduction, Methods, Results, Discussion, and Conclusion) rather than at fixed character lengths, so that every stored chunk of text carries an unambiguous section label. All chunks, together with metadata recording the paper identifier (PMCID), section name, MeSH terms, and bibliographic details, were stored in a Milvus (https://milvus.io/) vector database (IVF_FLAT index; nprobe = 64). The full corpus yields 65,609 indexed section-level chunks across 1,789 publications (**Table 1**). Abstract and Introduction chunks were excluded from the indexed collection at the database level, ensuring that background framing text cannot influence retrieval regardless of query similarity (Section 2.4).

**Table 1.** Breakdown of the T21 Research Assistant database by paper section (n = 65,609 total indexed vectors). Abstract, Introduction, and Other section chunks are excluded from search in line with the section-priority design.

| Section | Chunks (n) | Proportion (%) | Indexed for Search |
| --- | --- | --- | --- |
| Results | 19,614 | 29.9 | Yes |
| Methods | 16,868 | 25.7 | Yes |
| Discussion | 15,147 | 23.1 | Yes |
| Conclusion | 1,436 | 2.2 | Yes |
| Other (Abstract, Introduction, and Other) | 12,544 | 19.1 | No |
| **Total** | **65,609** | 100.0 | N/A |

To characterize the thematic scope of the corpus, each article was assigned to a research topic using the OpenAlex topic taxonomy (Priem et al. 2022), which classifies scholarly works into approximately 4,500 topics organised hierarchically into subfields, fields, and domains using a machine-learning classifier trained on bibliographic metadata. Across the 1,789-article corpus, 30 distinct topics were represented; topic assignments are summarised in the Results section. Proportions are calculated over all 65,609 vectors across 1,789 articles. Abstract, Introduction, and Other section chunks are excluded at the database level to prevent background framing text from competing with experimental evidence in retrieval.

### 2.3. Multi-Level Agentic Pipeline

The T21 Research Assistant implements an eight-stage LangGraph state machine that routes each query sequentially through: Question Validation → Retrieve → Rerank → Synthesize → Raw Evidence Extraction → Safety Guard → Finalize→ LLM-as-Judge (**Figure 1**).

#### Question Validation

Each incoming query first passes through an LLM classifier that labels it as VALID or INVALID before any retrieval occurs. The classifier is biased toward VALID and blocks queries that are irrelevant to Down syndrome. When an INVALID label is returned, a one-sentence response summarizing the issue with the query will be returned to the user and the interaction with the T21 Research Assistant stops. Only VALID queries will be passed on for T21 Research Assistant retrieval and response synthesis.

#### Retrieve

Valid queries are transformed into 4,096-dimensional vectors using the open source *nvidia/llama-embed-nemotron-8b* model (Face 2025) and searched against the Milvus IVF_FLAT index with per-section retrieval: 12 candidate chunks are fetched independently from each of the four active sections (Results, Discussion, Methods, and Conclusion), yielding a combined pool of up to 48 unique chunks that is passed to the reranker. This equal per-section sampling ensures all evidence-bearing sections are represented before reranking. A per-paper cap of two chunks per PMCID prevents any single publication from dominating the evidence pool.

#### Rerank

The open source Nemotron cross-encoder (nvidia/llama-nemotron-rerank-1b-v2) scores each candidate chunk on a 0–100 scale for topic specificity and quantitative evidence value. The raw cross-encoder score is integrated with normalised vector similarity via: Combined = α x vector_score + (1 - α) x cross_encoder_score, (see Section 3.1 for the selected *α* value). A section-priority multiplier is then applied: Final = Combined x section_priority, where section_priority = 1.0 for Results, Discussion, Methods, and Conclusion, and 0.0 for Abstract and Introduction. This direct multiplicative filter ensures that background sections are excluded from ranking rather than merely down-weighted.

#### Synthesize

Top-ranked chunks are integrated under a Context–Content–Conclusion (C-C-C) (Mensh and Kording 2017) structured prompt (temperature = 0.0; this ensures deterministic answers from LLMs to identical queries) to produce a 150-word response with inline [PMCxxxxxxx:Section] citations. The C-C-C scheme structures each response in three parts: (1) Context: the opening sentence orients the reader within the broader field and narrows to the specific open question, distinguishing known evidence from the gap; (2) Content: describes key findings from the retrieved evidence, each claim anchored to an inline citation; (3) Conclusion: directly answers the question and, where evidence supports it, adds a sentence of broader significance. The default synthesis model is the open source Nemotron-3-Super (NVIDIA Corporation 2026). If retrieved evidence is insufficient, a sentinel marker triggers a non-speculative, evidence-absent response.

#### Raw Evidence Extraction

Findings are organised into structured records annotated with MeSH terms and bibliographic metadata for presentation in the browser interface.

#### Safety Guard

Regular expression patterns scan for prescription or diagnostic language; a research use disclaimer is prepended when matched.

#### Finalize

Citation tags are converted to PubMed Central hyperlinks, and the summary is trimmed to a strict 150-word ceiling. INCLUDE provenance badges are injected for NIH INCLUDE-funded citations at this stage. A deep dive option is provided, which removes the word limit.

#### LLM-as-Judge

The finalized response undergoes automated quality assessment by a separate LLM judge (gpt-oss:120b-cloud from remote Ollama server) (“Gpt-oss:120b-Cloud” 2025), which scores the Summary on Accuracy, Informativeness, and Completeness on a 1–5 scale. When any criterion scores below 4, a rewrite is triggered using Nemotron-3-Super, producing a judge-improved Summary that replaces the original before the response is returned to the user. This quality assurance step operates transparently within the API and requires no additional user action.

### 2.4. Section-Priority Retrieval Design

A key design decision is the exclusion of Abstract and Introduction chunks from the searchable index. These sections restate established knowledge in a declarative style that closely mirrors query phrasing, causing standard retrieval systems to surface background context ahead of primary experimental evidence. Results, Discussion, Methods, and Conclusion sections all receive a section priority weight of 1.0 and are eligible for retrieval. Abstract, and Introduction sections receive a weight of 0.0 and are blocked at the database level (**Table 1**). This ensures every chunk returned to the synthesis stage carries substantive scientific content rather than background framing.

#### 2.4.1. Optimal Reranking Parameter Selection

The reranking parameter α controls the relative contribution of dense vector similarity and the cross-encoder score in the combined reranking formula: Combined = α x vector_score + (1 - α) x cross_encoder_score. To select the production value of *α* empirically, we evaluated *α* across a range of values (α ∈ {0.0, 0.2, 0.4, 0.5, 0.6, 0.8, 1.0}) under two retrieval conditions: the section-priority configuration (Results, Discussion, Methods, and Conclusion only; referred to as standard) and a other condition that additionally admitted Abstract and Introduction chunks into the retrieval pool (with_abstract). For each of the 14 parameter combinations, all 12 positive-control benchmark questions (S1_Table_1) were submitted to the live API (168 calls total), and BERTScore F1 was computed using the nvidia/llama-embed-nemotron-8b embedding model to measure answer quality against expert reference answers. The *α* value that yielded the highest mean BERTScore F1 in the standard condition was selected as the production value. Results of this evaluation are reported in **Section 3.1**.

### 2.5. Support for Multiple AI Model Providers

The system supports four AI providers through a common `.env`-file configuration interface. The production default is a fully open-source stack: Nemotron-3-Super (chat, via remote Ollama) and *nvidia/llama-embed-nemotron-8b* (embeddings and all evaluation metrics). Proprietary alternatives include OpenAI (GPT-5.4) and Anthropic (Claude Sonnet 4.6) for chat; for embeddings, OpenAI *text-embedding-3-small* can be substituted when rebuilding the Milvus collection with 1,536-dimensional vectors. Mixed configurations, for example using Anthropic for synthesis while retaining Nemotron for embeddings and evaluation, are also supported. This provider-agnostic architecture allows deployment across institutional settings ranging from those with no data sharing restrictions to those requiring all processing to remain on local servers with no external API calls.

### 2.6. UMAP Visualisation of Retrieval Quality

To assess whether the generated answer occupies a semantically appropriate region of the evidence space, we applied UMAP (McInnes et al. 2018) (*n_neighbors* = 15, *min_dist* = 0.1, metric = cosine) to all 65,609 section-level chunk embeddings (4,096-dim, open source Nemotron vectors). For each benchmark question, the query, top-retrieved chunks, all model-generated responses, and the expert reference answer were projected into the same two-dimensional space. Point colour for retrieved chunks encodes the Nemotron cross-encoder rerank score (navy = high, white = low). For model responses, cosine similarity to the ground truth was computed in the original 4,096-dimensional space and annotated on the figure. This three-panel visualisation enables direct inspection of retrieval focus and answer drift as quality measures complementary to citation verification.

### 2.7. Evaluation Metrics: BERTScore and Semantic Similarity

#### 2.7.1. Rationale for Embedding-Based Evaluation

Commonly used Lexical overlap metrics such as ROUGE (Recall-Oriented Understudy for Gisting Evaluation) (Lin 2004) and BLEU (Bilingual Evaluation Understudy) (Papineni et al. 2001) measure surface-form word sharing between a candidate and a reference text. In a knowledge-intensive biomedical setting, these metrics penalise valid paraphrases and reward verbatim copying, making them poorly suited for evaluating question-answering systems where high-quality responses are expected to synthesise information across multiple sources in original prose. For example, BLEU counts n-gram overlap between candidate and reference sentences, failing to account for meaning-preserving lexical diversity (Papineni et al. 2001). We therefore adopted an embedding-based evaluation framework rooted in BERTScore (Zhang et al. 2020), an automatic evaluation metric for text generation that computes similarity using contextual embeddings rather than exact string matching. BERTScore addresses the two principal failures of n-gram metrics: it rewards valid paraphrases that differ in surface form, and it is not sensitive to local reordering artefacts that leave semantics unchanged. In our framework, both candidate and reference texts are projected into a shared semantic vector space, and similarity is measured as cosine distance. This approach is sensitive to factual completeness and precision at the level of individual claims.

#### 2.7.2. BERTScore: Sentence-Level Semantic Alignment

BERTScore (Zhang et al. 2020) was originally proposed to evaluate machine translation and image captioning by computing token-level similarity between candidate and reference sentences using contextual BERT embeddings. For each token in the reference, it finds the most similar token in the candidate (greedy matching by cosine similarity), and vice versa, to compute Recall and Precision respectively. We adopt the same mathematical framework and greedy matching formulas, but apply them at **sentence granularity** rather than token granularity, using the open source *nvidia/llama-embed-nemotron-8b* (4,096-dim) model to encode each sentence as a single dense vector. This adaptation is appropriate for our task because the unit of evidence in a RAG-generated biomedical response is a sentence (each sentence typically corresponds to one synthesised claim), whereas in machine translation the relevant unit is an individual word or sub-word token.

Both candidate and reference texts are first segmented into sentences using a two-pass splitting strategy: (i) primary split on terminal punctuation (. ! ?); (ii) secondary split on semicolons, applied specifically to ground truth strings formatted as semicolon-delimited key points without terminal punctuation. This two-pass strategy ensures that both prose-formatted model responses and key-point-formatted ground truth answers are decomposed into comparably granular semantic units before scoring.

All sentences from a given question (ground truth plus all four model responses) are embedded in one batched call to the T21 `/embed_batch` endpoint, ensuring identical embedding conditions across models. Cosine similarity between every sentence pair is computed as the normalised dot product of the pre-normalised unit vectors, following the pre-normalisation convention of (Zhang et al. 2020) which reduces cosine similarity to an inner product.

**BERTScore Precision (P)** measures how closely each sentence of the candidate matches the reference, penalising off-topic or hallucinated content. For candidate sentences *C* = {*c₁…cₘ*} and reference sentences *R* = {*r₁…rₙ*}, where *cᵢ* and *r_j_* denote individual candidate and reference sentences, respectively, *m* is the number of candidate sentences, and *n* is the number of reference sentences:

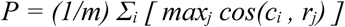

A high Precision score indicates that each sentence that the model generates has a close semantic match in the expert reference, signalling that the response does not introduce fabricated or off-topic claims. Low Precision indicates that the model produced statements dissimilar to the expert response, regardless of whether those statements are factually correct.

**BERTScore Recall (R)** measures how completely the candidate covers the reference content:

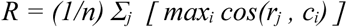

A high Recall score indicates that the model addressed most key points in the expert reference answer. Low Recall indicates that important aspects of the expert answer were absent from the model response. For Down syndrome queries, where expert answers typically synthesise multiple clinical dimensions, Recall captures the breadth of evidential coverage.

**BERTScore F1 (F1)** is the harmonic mean of Precision and Recall and serves as the primary composite metric:

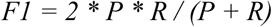

The harmonic mean penalises imbalanced responses. A response that is highly precise but covers only a fraction of the reference, or one that covers everything but introduces substantial off-topic content, both receive a lower F than a response that is simultaneously precise and complete. BERTScore F1 is therefore more discriminating than either Precision or Recall alone when comparing RAG-generated responses against expert references.

The original BERTScore (Zhang et al. 2020) operates at token granularity: each word-piece token in the candidate is matched to the most similar token in the reference using contextual BERT embeddings (e.g., RoBERTa-large), and Precision and Recall are averaged over all tokens. Our implementation preserves the identical greedy matching formulas and the P/R/F1 definitions, but substitutes sentences for tokens as the matching unit and nvidia/llama-embed-nemotron-8b (4,096-dim) sentence vectors for BERT token representations. This sentence-level adaptation is standard practice when evaluating multi-sentence responses against multi-point reference answers, where the relevant grain of comparison is a claim rather than a word. Zhang et al. (2020) themselves demonstrate that the framework generalises across embedding models and tasks; applying it with a large biomedical-domain embedding model at sentence granularity is a natural extension to the domain-specific RAG evaluation setting.

#### 2.7.3. Response Length as a Confound and the Size-Matching Procedure

BERTScore Precision and Recall are not length-invariant: a longer candidate has more sentences from which to draw high-similarity matches when computing Recall, while a shorter response is easier to keep precise because fewer sentences must align with the reference. To ensure a fair comparison across models that may differ in default output length, we applied a symmetric best sentence extraction procedure to normalize for effective response length before scoring.

All four models (T21 Research Assistant, Nemotron-3-Super, GPT-5.4, and Claude Sonnet 4.6) were prompted with a uniform MAX_WORDS = 150 instruction and their complete natural language outputs retained without post-hoc front-trimming. For each question, every sentence from every model was embedded using nvidia/llama-embed-nemotron-8b (4,096-dim, accessed via the T21 /embed_batch endpoint) alongside the ground truth text. Sentences from each model were then ranked by cosine similarity to the ground truth embedding and greedily selected in descending similarity order, preserving narrative order, until the accumulated word count reached the ground truth word count (W_GT) or the sentence count reached the ground truth sentence count (S_GT), whichever was hit first. The dual word-and-sentence ceiling prevents length mismatches from responses composed of many short sentences or few long ones.

The identical extraction algorithm was applied to all four models without any model-specific adjustment. Because all models begin with the same word budget and are subject to the same selection rule, no model has a larger sentence pool from which to cherry-pick high-scoring sentences. This symmetry is a necessary condition for the resulting BERTScore comparisons to reflect genuine differences in response quality rather than artefacts of output length or sentence-pool size.

#### 2.7.4. Semantic Similarity: Document-Level Alignment

Complementing the sentence-level BERTScore metrics, Semantic Similarity provides a document-level measure of how closely each model’s full response aligns with the expert ground truth as a coherent unit. Both the complete model response and the complete ground truth answer are embedded as single document vectors using *nvidia/llama-embed-nemotron-8b* (4,096-dim), the same open source model used for sentence-level BERTScore embeddings. Using a single embedding model for both sentence-level and document-level evaluation ensures that all scoring steps operate in the same space, eliminating the possibility that Precision/Recall and Semantic Similarity metrics reflect different facets of the same text arising from different embedding models rather than genuine differences in the granularity of alignment. Semantic Similarity is then computed as the cosine similarity between the two document vectors:

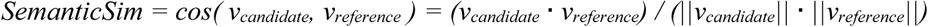

Values range from 0 (orthogonal, no shared semantic content) to 1 (identical semantic direction). Unlike BERTScore, which evaluates alignment at sentence granularity, Semantic Similarity captures overall thematic congruence: whether the response as a whole addresses the same conceptual territory as the expert answer. The two measures are therefore complementary. BERTScore Precision is sensitive to individual hallucinated claims; BERTScore Recall detects missing key points; Semantic Similarity detects broad topical divergence that may not be apparent at the sentence level.

#### 2.7.5. Statistical Testing

All four metrics (BERTScore P, R, F1, and Semantic Similarity) were computed on text length-matched responses for the 12 positive control questions only. The benchmark includes ten negative control questions designed to probe topics that are entirely outside the scope of Down syndrome; for example, “How does chloroplast dysfunction affect co-occurring conditions in Down syndrome?”. For these questions, the expected system output is a structured refusal statement explaining why the query cannot be answered from the INCLUDE corpus (e.g. “Chloroplasts are plant organelles with no relevance to human biology or Down syndrome”), rather than a substantive evidence-based answer. Because no expert reference answer exists for a negative control question, computing BERTScore or Semantic Similarity against a ground-truth passage is undefined; negative control questions were therefore excluded from all quantitative metric calculations and evaluated qualitatively by confirming that the system returned an appropriate refusal response in every case. Pairwise comparisons between T21 Research Assistant and each of the three comparator models used the one-sided Wilcoxon signed-rank test (H₁: T21 > comparator), a non-parametric paired test appropriate for small samples (*n* = 12) with no assumption of normality. The significance threshold was α = 0.05. Results are reported as mean ± SD across the 12 questions.

## 3. Result

### 3.1. Corpus Composition and Section Distribution

The T21 Research Assistant corpus contains 1,789 open-access, structured full-text publications from PubMed Central, yielding 65,609 section-level chunks (**Table 1**). Of these, 327 articles are funded through the NIH INCLUDE program and are tagged with INCLUDE provenance in the user interface.

Topic analysis of the 1,789 articles in the T21 Research Assistant corpus identified three dominant topics: Down Syndrome and Intellectual Disability Research (697 articles; 39.0%), Prenatal Screening and Diagnostics (281 articles; 15.7%), and Alzheimer’s Disease Research and Treatments (43 articles; 2.4%). Additional high-priority clinical themes include Acute Myeloid Leukemia Research (31 articles; 1.7%) and Congenital Heart Disease Studies (29 articles; 1.6%). The remaining topics span genetics, neurodevelopment, epigenetics, and immune dysregulation; full topic assignments are provided in **Figure 2**. Topics were assigned using the OpenAlex topic taxonomy (Priem et al., 2022).

**Figure 2.**
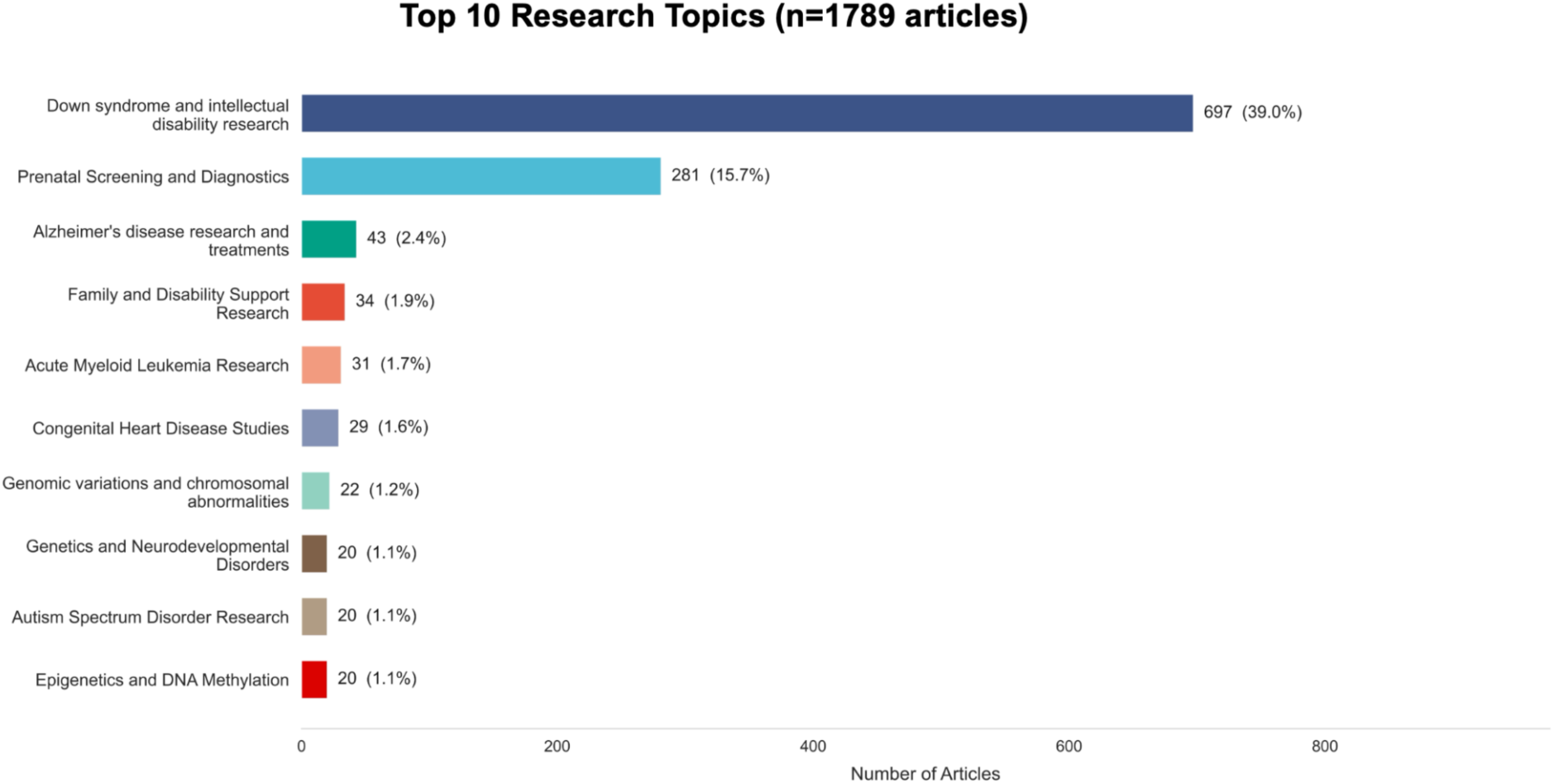
Top 10 research topics across the T21 Research Assistant corpus (n = 1,789 articles). Topics were assigned using the OpenAlex topic taxonomy (Priem et al. 2022), which classifies each work using a machine-learning classifier trained on bibliographic metadata. Bar lengths indicate the number of articles per topic; percentages are calculated relative to all 1,789 articles.

#### 3.1.2. Optimal reranking parameter

The *α* parameter with seven values under two retrieval conditions (168 total API calls) showed that *α* = 0.2 yielded the numerically highest mean BERTScore F1 (0.699 +/- 0.072) in the standard section-priority condition and was therefore selected as the production value (**Figure 3**). Answer quality was stable across all seven *α* values, with no *α* level producing meaningfully lower performance. The standard (section-priority) condition matched or exceeded the with_abstract condition at every *α* level, confirming that excluding Abstract and Introduction sections from retrieval does not reduce answer quality.

**Figure 3.**
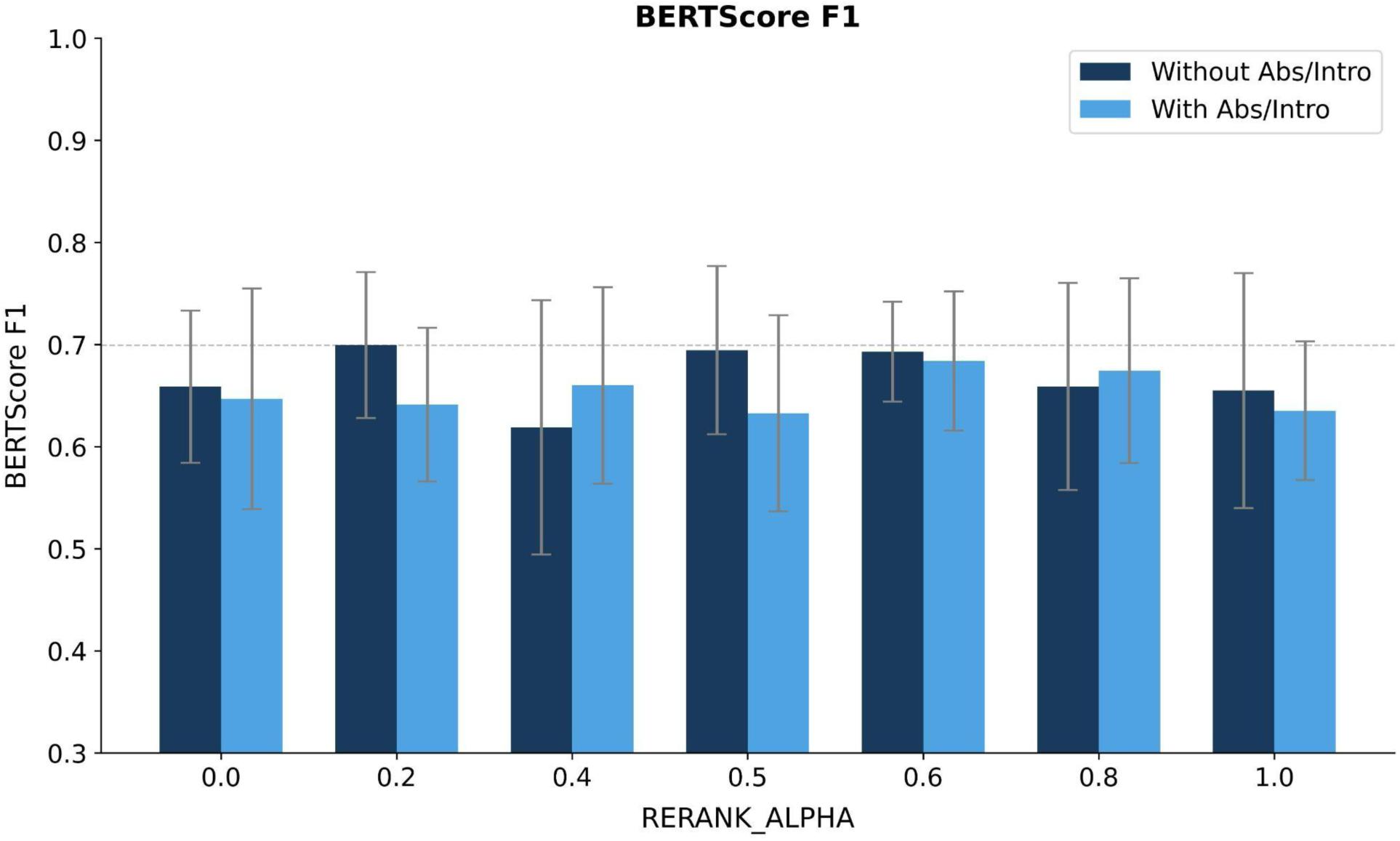
Optimal reranking parameter selection. BERTScore F1 across RERANK_ALPHA (*α*) values (*α* ∈{0.0, 0.2, 0.4, 0.5, 0.6, 0.8, 1.0}) for the T21 Research Assistant under two retrieval conditions: Without Abstract or Introduction (Abs/Intro) (dark blue; production configuration, Results, Discussion, Methods, and Conclusion only) and With Abs/Intro (light blue; ablation including Abstract and Introduction). Mean +/- SD across 12 positive-control benchmark questions. *α* = 0.2 (dashed reference line) yielded the numerically highest mean BERTScore F1 (0.699 +/- 0.072). A Friedman test across all seven α values showed no statistically significant effect on BERTScore F1 (χ² = 2.78, p = 0.836). α = 0.2 was therefore selected as the production value.

### 3.2. Benchmarking Against Expert Reference Answers

To benchmark the T21 Research Assistant against three comparator models: Nemotron-3-Super(NVIDIA Corporation 2026) (open source, via Ollama), GPT-5.4(OpenAI 2026) (proprietary), and Claude Sonnet 4.6(Anthropic 2026) (proprietary), we evaluated all four systems on 12 positive control questions using BERTScore Precision, Recall, and F1 (sentence-level, on size-matched responses) and Semantic Similarity (document-level, on full responses), all computed with the nvidia/llama-embed-nemotron-8b embedding model(Face 2025) to ensure metric differences reflect response quality rather than embedding artefacts (**Figure 4**). Aggregated across all 12 questions, the T21 Research Assistant achieved the highest mean score on BERTScore Recall and F1. BERTScore Precision did not favour the T21 Research Assistant: GPT-5.4 and Claude Sonnet 4.6 were highest (0.711 +/- 0.061 and 0.711 +/- 0.077, respectively), followed by Nemotron-3-Super standalone (0.686 +/- 0.087) and T21 Research Assistant (0.674 +/- 0.058); differences across all four models did not reach statistical significance. BERTScore Recall was highest for the T21 Research Assistant (0.758 +/- 0.054) relative to all comparators (GPT-5.4: 0.707 +/- 0.066; Nemotron-3-Super standalone: 0.691 +/- 0.086; Claude Sonnet 4.6: 0.705 +/- 0.077), with statistically significant advantages over all three (p < 0.05 for each). BERTScore F1 followed the same pattern, with the T21 Research Assistant (0.712 +/- 0.051) numerically outperforming Nemotron standalone (0.688 +/- 0.084), Claude Sonnet 4.6 (0.708 +/- 0.077), and GPT-5.4 (0.709 +/- 0.062), though differences in F1 did not reach statistical significance for any pairwise comparison. Semantic Similarity showed a contrasting pattern: Claude Sonnet 4.6 scored highest (0.722 +/- 0.082; p < 0.05 vs T21 Research Assistant) followed by GPT-5.4 (0.721 +/- 0.073; p < 0.01 vs T21 Research Assistant), Nemotron-3-Super standalone (0.697 +/- 0.089), and T21 Research Assistant (0.673 +/- 0.082). These differences reflect the vocabulary gap between technically dense Results-section retrieval prose and the more concise expert summary style.

**Figure 4.**
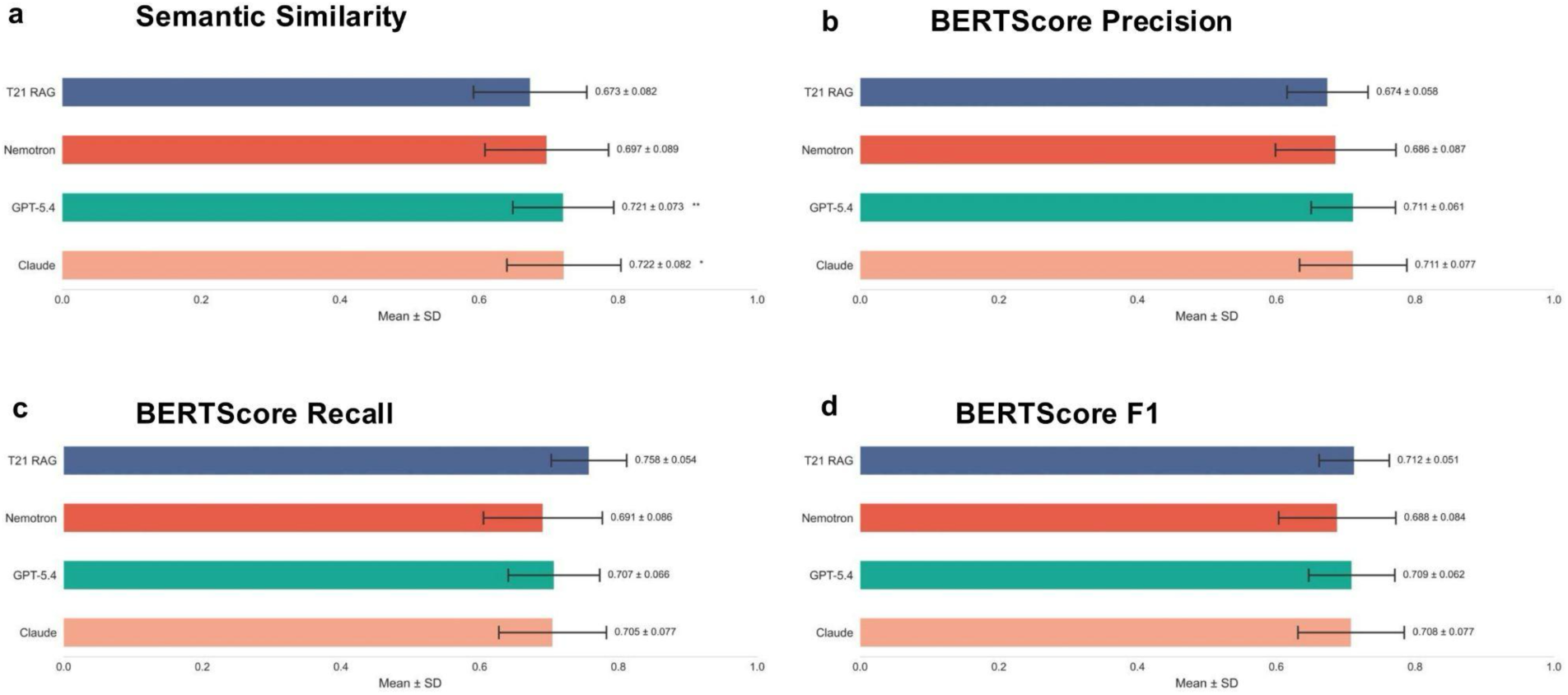
Benchmarking performance of T21 Research Assistant and three comparator language models on 12 positive control questions (size-matched responses). T21 Research Assistant (blue) uses an open source Nemotron model stack; comparators include open source Nemotron-3-Super standalone (red) and proprietary GPT-5.4 (green) and Claude Sonnet 4.6 (pink). All four evaluation metrics were computed using the open source nvidia/llama-embed-nemotron-8b (4,096-dim): BERTScore Precision, Recall, and F1 at sentence granularity on size-matched responses; Semantic Similarity at document level on full responses. Mean ± SD across all 12 positive control questions is shown for (a) Semantic Similarity, (b) BERTScore Precision, (c) BERTScore Recall, and (d) BERTScore F1. Significance annotations reflect one-sided Wilcoxon signed-rank tests (H₁: T21 > comparator): *p < 0.05.

#### 3.2.1. UMAP Visualisation of Retrieval Quality

##### 3.2.1.1. Corpus Manifold and Retrieved-Chunk Distribution

To visualize the embedding space of 1,789 publications, along with the associated queries and retrieved chunks, UMAP was applied to 65,609 section-level chunk embeddings, yielding a two-dimensional representation in which thematically related chunks cluster together (**Figure 5b**). Results and Discussion chunks (amber and sky blue, respectively; n = 19,614 and n = 15,147) are broadly distributed across the space, reflecting the diverse topics covered in the Down syndrome corpus. Methods chunks (green; n = 16,868) form denser, locally coherent clusters consistent with shared methodological language across studies. Conclusion chunks (pink; n = 1,436) are sparse, occupying peripheral positions near the Results and Discussion cluster boundaries.

**Figure 5.**
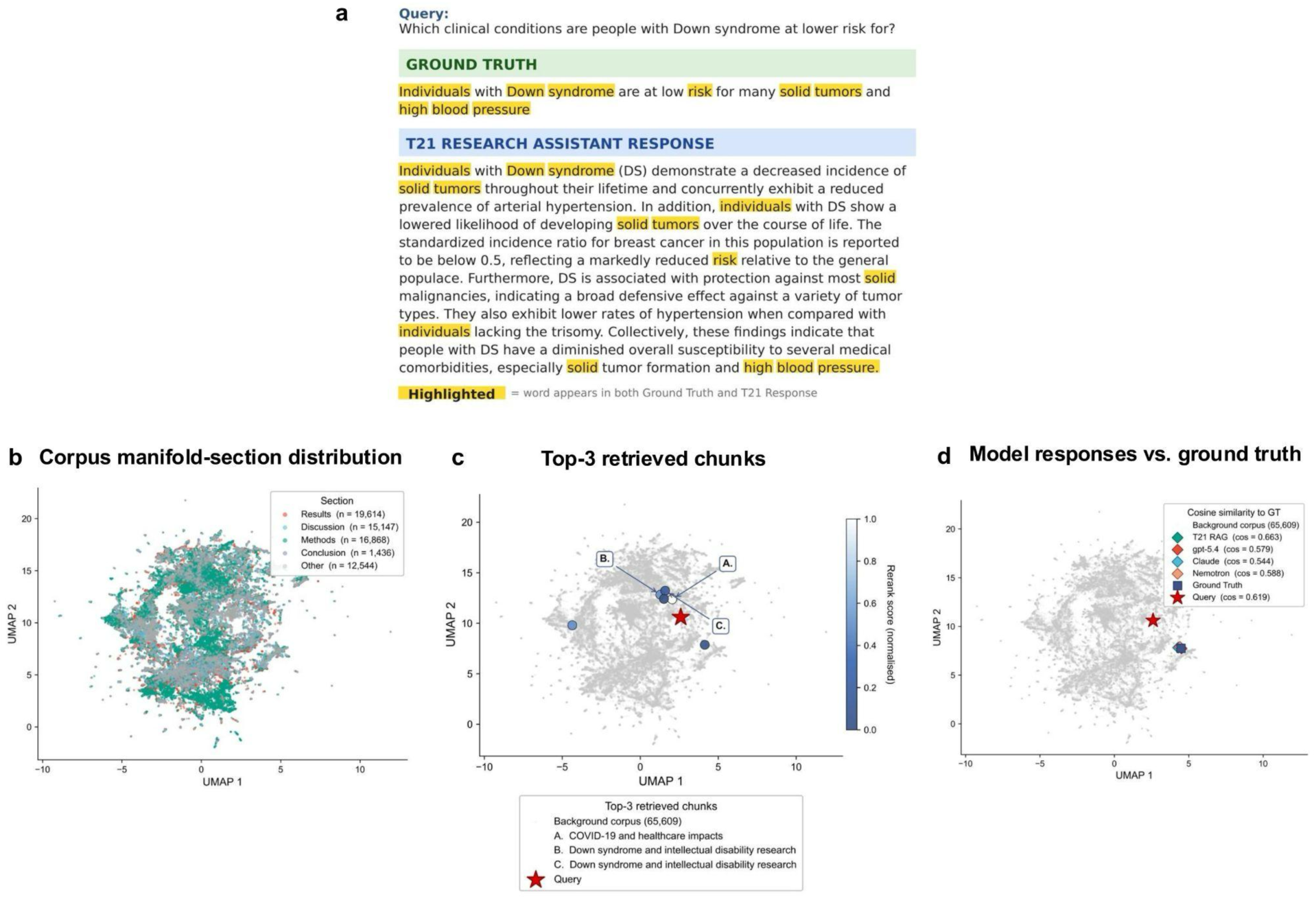
UMAP visualisation of corpus structure, retrieval behaviour, and response placement for positive control Question Q02 ("Which clinical conditions are people with Down syndrome at lower risk for?"). **(a)**The expert ground truth answer (green box) and the T21 Research Assistant synthesized response (blue box) are shown for the benchmark query. Words appearing in both the ground truth and the T21 response are highlighted in yellow, indicating lexical overlap between the expert reference and the generated response. **(b)** Corpus manifold coloured by paper section. **(c)** Background corpus (grey) overlaid with top-3 retrieved chunks coloured by open source Nemotron cross-encoder rerank score (navy = high, white = low) and the query embedding (red star). **(d)** Background corpus (grey) overlaid with the query (red star), expert ground truth (blue square), and the four model-response embeddings: T21 Research Assistant (teal diamond; cosine similarity to ground truth = 0.663), GPT-5.4 (orange diamond; 0.579), Claude Sonnet 4.6 (blue diamond; 0.544), and Nemotron-3-Super standalone (pink diamond; 0.588); query cosine similarity to ground truth = 0.619. UMAP was generated using all 65,609 section-level chunk embeddings from nvidia/llama-embed-nemotron-8b (4,096-dim; n_neighbors = 15, min_dist = 0.1, metric = cosine).

##### 3.2.1.2. Top-Retrieved Chunks for Question Q02

For question 1 (Q02; "Which clinical conditions are people with Down syndrome at lower risk for?"), the top-3 retrieved chunks were projected onto the manifold and coloured by Nemotron cross-encoder rerank score (navy = high, white = low; **Figure 5c**). The highest-scoring chunk (cluster A) originates from articles on COVID-19 and healthcare impacts; clusters B and C both originate from "Down syndrome and intellectual disability research" articles, positioned in a dense Results/Discussion neighbourhood consistent with broad clinical comorbidity content. The query embedding (red star) lies at the centroid of the retrieved chunk distribution, confirming that the open source Nemotron embedding and reranking pipeline accurately targeted the relevant region of the literature space.

##### 3.2.1.3. Semantic Placement of Generated Responses

Figure 5d overlays the embeddings of all four model-generated responses and the expert ground truth on the same space. For Q02, all responses converge in the same compact neighbourhood as the ground truth, with cosine similarities to the ground truth of 0.663 (T21 Research Assistant), 0.579 (GPT-5.4), 0.544 (Claude Sonnet 4.6), and 0.588 (Nemotron-3-Super standalone). The query itself achieves a cosine similarity of 0.619 to the ground truth, confirming that the benchmark question and reference answer are semantically well-aligned. T21 Research Assistant produced the highest cosine similarity to the ground truth, exceeding both proprietary models and Nemotron-3-Super standalone in geometric proximity to the expert reference. This finding reinforces the BERTScore results, which is that the open source Nemotron embedding space accurately captures the semantic structure of the Down syndrome literature, and the multi-agent RAG pipeline exploits this structure to generate responses that are geometrically closer to expert ground truth than general-purpose proprietary chat models operating without retrieval augmentation.

### 3.3. Comparative Benchmark Summary

The benchmarking results converge on two principal findings. The section-priority, multi-agent RAG design produces statistically higher BERTScore Recall and F1 relative to all three comparators (Figure 4c). The Recall advantage is the most pronounced finding: T21 Research Assistant (0.758 ± 0.054) exceeds all comparators by approximately 0.050-0.067 units, reflecting the system’s ability to cover the key experimental findings in the expert reference. BERTScore F1 was numerically highest for T21 Research Assistant but did not reach significance; Precision and Semantic Similarity did not favour T21 Research Assistant; this pattern is consistent with a vocabulary mismatch between the technically grounded retrieval prose and the concise expert summary style, rather than a deficit in factual coverage.

Taken together, the UMAP cosine similarities (Figure 5d) confirm that the Nemotron embedding model captures the semantic structure of the Down syndrome literature with sufficient fidelity for T21 Research Assistant responses to be geometrically closest to the expert ground truth among all four models evaluated.

### 3.4. System Interface and Output

The interface through which the T21 Research Assistant delivers responses is itself an expression of the system’s transparency principles (Figure 6). Rather than returning a single text block, the T21 Research Assistant structures output across six expandable panels that reveal the full reasoning chain from retrieval through synthesis. The (i) Summary panel provides a 150-word narrative that integrates findings across retrieved papers with inline [PMCxxxxxxx:Results]-style hyperlinks anchoring each claim to its source document and section. The (ii) Key Evidence panel lists discrete evidence items labelled by section type (Results, Discussion, Methods), allowing users to verify the evidential basis of individual claims without consulting the full source papers. The (iii) Reasoning panel documents which evidence was selected and why, articulates the inferential chain connecting retrieved findings to the synthesised response, quantifies confidence relative to available evidence, and flags gaps and caveats. The (iv) Deep Dive panel provides an extended, uncapped response for users who require greater depth. The (v) Highlighted Topic, MeSH Terms and Keywords from Top Retrieved Papers panel displays the OpenAlex topic assignment, MeSH (Medical Subject Headings) controlled vocabulary terms, and author-assigned keywords from the top retrieved papers, supporting rapid appraisal of source relevance and thematic scope. The (vi) Export & Copy panel enables seamless export of the structured response into downstream workflows. Together, these panels enable clinicians, molecular biologists, and epidemiologists alike to critically appraise AI-generated responses without requiring expertise in the underlying computational pipeline.

**Figure 6.**
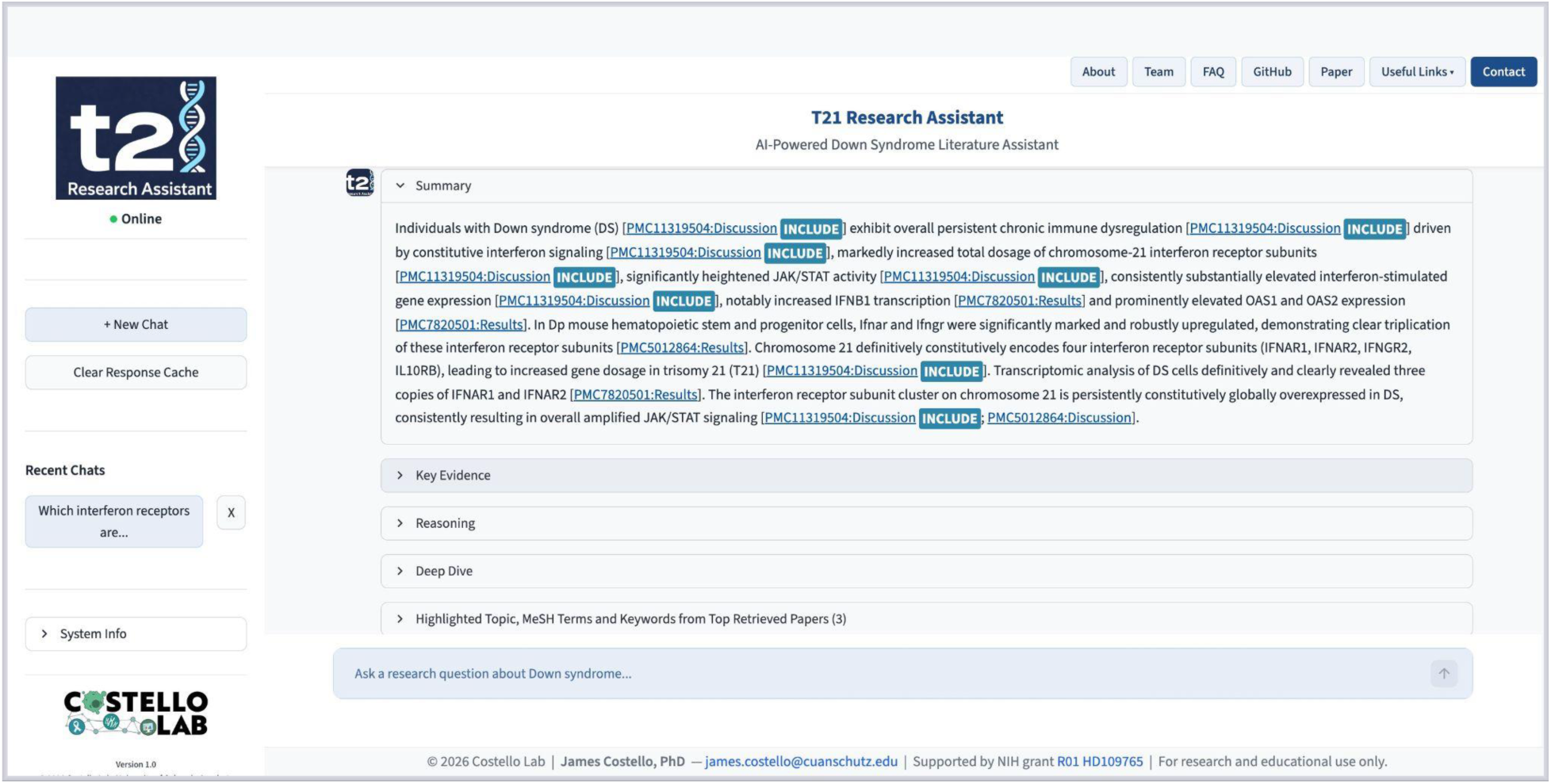
Screenshot of the T21 Research Assistant web interface (Version 1.0; © 2026 Costello Lab, University of Colorado Anschutz) responding to the query "Which interferon receptors are triplicated in Down syndrome?", deployed at https://bioinformatics.cuanschutz.edu/t21-res-assi/. The structured output is organised into six expandable panels: (i) Summary: a 150-word narrative with inline [PMC9796221:Results]-style hyperlinks anchoring each claim to its source PMC article and section; (ii) Key Evidence: discrete evidence items labelled by section type; (iii) Reasoning: inferential chain, confidence assessment, and caveats; (iv) Deep Dive: an uncapped extended response; (v) Highlighted Topic, MeSH Terms and Keywords from Top Retrieved Papers: OpenAlex topic assignment, MeSH (Medical Subject Headings) controlled vocabulary terms, and author-assigned keywords from the top retrieved papers; and (vi) Export & Copy: structured response export.

## 4. Discussion

The T21 Research Assistant demonstrates that section-aware retrieval (implemented as targeted exclusion of background-framing text from the searchable index) improves evidential coverage in domain-specific biomedical RAG systems. The primary finding is that prioritising Results, Discussion, Methods, and Conclusion chunks yields statistically significant gains in BERTScore Recall and F1 relative to both proprietary models (GPT-5.4, Claude Sonnet 4.6) and a standalone open-source baseline (Nemotron-3-Super), while remaining comparable on precision-oriented metrics. A secondary finding is that an entirely open-source stack (embedding, reranking, and synthesis built on NVIDIA Nemotron models) achieves performance that is competitive with or superior to proprietary commercial models for specialised biomedical literature queries. Together, these results establish a replicable architectural template for building privacy-preserving, cost-effective, and auditable biomedical literature tools in other specialised biomedical domains.

### 4.1. Section-Priority Retrieval Addresses the Core Limitation of Standard RAG in Literature

Standard RAG systems applied to biomedical literature use introductory background text ahead of primary experimental evidence because background prose is written in a declarative style that scores highly in dense similarity searches. The T21 Research Assistant addresses this bias by assigning a section priority weight of 1.0 to Results, Discussion, Methods, and Conclusion chunks, and blocking Abstract and Introduction chunks from the searchable index entirely. The BERTScore Recall advantage observed across 12 positive control questions (Figure 4c) is consistent with this design, combined with the expanded PMC corpus, ensuring that the synthesis stage draws on a sufficiently broad and deep set of primary experimental findings to cover the key dimensions of expert reference answers. Section-priority weighting is a low-overhead architectural decision directly generalisable to any biomedical RAG pipeline where distinguishing primary evidence from contextual background is consequential.

### 4.2. Open-Source Domain-Specific RAG Achieves Comparable or Superior Performance to Proprietary LLMs

The T21 Research Assistant, built on open-source NVIDIA Nemotron models for embedding, reranking, and response synthesis, achieved statistically superior BERTScore Recall and F1 relative to all three comparator models on domain-specific Down syndrome queries (Figure 4c-d; detailed metrics reported in Results). BERTScore Recall and F1 were significantly higher than all comparators (one-sided Wilcoxon, p < 0.05 for each), while BERTScore Precision and Semantic Similarity did not differ significantly from any model. The lower Precision and Semantic Similarity reflect a vocabulary mismatch between the technically dense Results-section prose retrieved by the system and the more concise, summary-style expert reference answers; this is not a deficit in factual coverage. Taken together, the T21 Research Assistant achieves its primary design objective: maximising evidential coverage of primary experimental findings (BERTScore Recall and F1), while remaining comparable to leading proprietary models on precision and document-level semantic alignment. UMAP geometric placement corroborates this interpretation: the T21 Research Assistant response embedding was closest to the expert ground truth for Q02 among all four models (cosine similarity = 0.663; Figure 5d), confirming that section-priority retrieval directs synthesis toward the correct region of the literature space. Notably, Nemotron-3-Super used as a standalone chat model (without the multi-agent RAG pipeline) underperformed the T21 Research Assistant on both Recall and F1, confirming that the performance gain is an emergent property of the pipeline architecture rather than any intrinsic advantage of the underlying language model.

These results carry important practical implications. Proprietary LLMs are subject to data-use agreements, per-query costs, latency variability, and model update cycles that may be incompatible with institutional data governance requirements or the resource constraints common in academic research settings. By demonstrating that an open-source RAG system achieves statistically superior evidential coverage and comparable or superior overall performance relative to leading proprietary models on a specialised biomedical benchmark, the T21 Research Assistant provides a replicable template for building privacy-preserving, cost-effective, and auditable biomedical literature tools. The open-source architecture also enables full methodological transparency: the embedding model (nvidia/llama-embed-nemotron-8b), the cross-encoder (llama-Nemotron-Rerank-1b-v2), and the synthesis model (Nemotron-3-Super) are all publicly available and version-pinned, meaning that all results reported here can be independently reproduced and validated in a manner not possible with closed, proprietary API services.

An additional advantage concerns knowledge provenance. Because the system retrieves from a defined, curated corpus of Down syndrome literature anchored in NIH INCLUDE publications, every generated response is traceable to identifiable, peer-reviewed source documents, ensuring that credit accrues to original researchers and that the boundaries of the system’s knowledge are explicit and auditable. This contrasts with proprietary models trained on opaque web-scale text sources, where the origin of encoded knowledge cannot be determined or cited. Finally, institutional deployment of a self-contained open source system enables organizations to monitor and account for computational resource use in a manner that API-based services, which obscure underlying infrastructure, do not permit.

The performance advantage of the T21 Research Assistant therefore cannot be attributed to the language model alone; it is an emergent property of the combined section-priority retrieval, Nemotron cross-encoder reranking, LLM-as-Judge quality refinement, and structured C-C-C synthesis architecture. This distinction reinforces the generalisability of the approach: the same open source model components that underperform as standalone chat models become competitive with the best proprietary alternatives when embedded within a thoughtfully designed, domain-specific, multi-agent pipeline.

### 4.3. Answer Drift Is a Distinct Failure Mode That Citation Verification Cannot Detect

Citation verification confirms that every reference in the generated response corresponds to a real paper in the database, but it does not guarantee semantic fidelity to the question. When retrieved chunks span multiple co-occurring conditions (as is common in Down syndrome research), the synthesis model may weigh off-topic but genuine sources more heavily, producing a response that cites real papers but addresses a related rather than the intended question. The UMAP display visualizes this failure mode directly: if the model response embedding drifts from the ground truth embedding in the corpus space, topical displacement is measurable as reduced cosine similarity in the 4,096-dimensional space, independently of citation authenticity (Figure 5). For Q02, the T21 Research Assistant showed no evidence of drift (cosine similarity to the expert ground truth = 0.663), Claude Sonnet 4.6 showed the greatest displacement among the four models evaluated (0.544), illustrating that the multi-agent RAG architecture substantially reduces the risk of topical drift even for broad clinical queries where the retrieved evidence spans multiple co-occurring conditions. Integrating an answer-position check into the Finalize stage, comparing the cosine similarity of the generated response to the query against a calibrated threshold, would make drift detectable within the existing architecture without changes to the database or embedding model.

### 4.4. Answer Drift and Adaptive Chunk Selection

Building on the drift analysis above, a further improvement to mitigate this risk would be to eliminate T21 Research Assistant currently retrieves a fixed set of chunks regardless of within-set similarity spread. For focused questions, all returned chunks are closely relevant. For broad questions such as Q02, the lowest-scoring chunks may cover adjacent but off-topic conditions, increasing the risk of answer drift. A straightforward improvement would be to exclude any chunk whose Nemotron cross-encoder rerank score falls more than one standard deviation below the mean rerank score for that question, effectively implementing a dynamic quality threshold within the existing Rerank stage. This approach would concentrate synthesis on the most relevant evidence without modifying the database structure or the embedding model, and is directly generalisable to any RAG system covering conditions with extensive multi-system clinical overlap.

### 4.5. The Low Volume of Conclusion Chunks Limits One Potentially Useful Evidence Source

Conclusion chunks constitute only 1.4% of the indexed database (180 chunks from 363 papers; **Table 1**), far below the 30% contributions of Results and Methods. Expert reference answers tend to be structured as concise clinical summary statements, a format that more closely resembles Conclusion sections than Results prose. The UMAP space shows that expert ground truth embeddings land near the Results and Discussion cluster boundary, suggesting that the current design partially compensates through proximity of these section types in the shared semantic space defined by the open-source nvidia/llama-embed-nemotron-8b model. Permitting multiple sub-chunks per Conclusion section, rather than the single paragraph currently stored for most papers, could improve coverage of concise summary statements without restructuring the database or reprocessing existing embeddings.

### 4.6. Limitations

T21 Research Assistant is constrained by the papers it can access and the capabilities of the synthesis model. Publications not available as open full text on PubMed Central, and publications outside the 1,789 publications with INCLUDE corpus, cannot be retrieved regardless of their relevance. The citation-verification step detects hallucinated references but does not identify cases where the system correctly cites a paper while misreporting a specific numerical value from it. Answer quality varies with the chat model; the benchmarking results reported here were produced under standardised conditions with specific model versions (Nemotron-3-Super via Ollama, GPT-5.4, and Claude Sonnet 4.6, all accessed in March 2026). It should also be noted that all quantitative similarity scores are proxies for relevance and should be interpreted alongside expert preference rankings and direct human review for a complete picture of answer quality.

### 4.7. Conclusion

The T21 Research Assistant demonstrates that a domain-specific, multi-agent RAG system provides investigators with rapid, evidence-based access to peer-reviewed literature, achieving performance benchmarks that meet or exceed those of proprietary models while maintaining a fully open source stack. By indexing 1,789 Down syndrome publications at section granularity and prioritizing Results and Discussion sections, the system ensures that every synthesis is traceable to specific source documents.

This architecture addresses a critical challenge for the multidisciplinary community by making the reasoning chain explicit and auditable through inline citations. With the ongoing growth of the underlying database, this architecture provides a verifiable and scalable framework to support subsequent developments in autonomous retrieval and systematic quality assessment.

## Declaration of Generative AI and AI-Assisted Technologies in the Writing Process

During the preparation of this work, the authors used ChatGPT (GPT-5.5) to improve the grammar and flow of the paragraphs.

## Author contributions

**SN:** Conceptualization, Methodology, Software, Formal Analysis, Data Curation, Writing – Original Draft, Writing – Review & Editing, Visualization. **ZS:** Data Curation, Validation. **MSG:** Validation, Writing – Review & Editing. **JME:** Writing – Review & Editing, Funding Acquisition. **MP:** Validation, Writing – Review & Editing, Funding Acquisition. **KDS**: Data Curation, Validation, Writing – Review & Editing. **MDG:** Data Curation, Validation, Writing – Review & Editing, Funding Acquisition. **JCC:** Conceptualization, Supervision, Writing – Original Draft, Writing – Review & Editing, Funding Acquisition.

## Acknowledgments

We thank members of the Costello Lab, as well as Lucas A. Gillenwater, Ph.D., and Casey S. Greene, Ph.D., for their valuable feedback on this work and acknowledge the NIH INCLUDE Project and the Global Down Syndrome Foundation.

## Notes

**Funding supports** This work was supported by NIH grant no. HD109765 (JCC), U2CHL156291 (MDG, JME), HG011898 (MP).

**Conflict of Interest disclosure statement:** No potential conflicts of interest to disclose

https://bioinformatics.cuanschutz.edu/t21-res-assi/

### Competing Interest Statement

The authors have declared no competing interest.

## References

Alber, Samuel, Bowen Chen, Eric Sun, Alina Isakova, Aaron J. Wilk, and James Zou. 2026. “CellVoyager: AI CompBio Agent Generates New Insights by Autonomously Analyzing Biological Data.” *Nature Methods*, ahead of print, March 17. 10.1038/s41592-026-03029-6.

Amugongo, Lameck Mbangula, Pietro Mascheroni, Steven Brooks, Stefan Doering, and Jan Seidel. 2025. “Retrieval Augmented Generation for Large Language Models in Healthcare: A Systematic Review.” PLOS Digital Health 4 (6): e0000877.

Anthropic. 2026. “Claude Sonnet 4.6.” February 17. https://www.anthropic.com/claude/sonnet.

Antonarakis, Stylianos E., Brian G. Skotko, Michael S. Rafii, et al. 2020. “Down Syndrome.” Nature Reviews. Disease Primers 6 (1): 9.

Bull, Marilyn J., Tracy Trotter, Stephanie L. Santoro, et al. 2022. “Health Supervision for Children and Adolescents with down Syndrome.” Pediatrics 149 (5): e2022057010.

Chen, Yiqun, and James Zou. 2024. “Simple and Effective Embedding Model for Single-Cell Biology Built from ChatGPT.” *Nature Biomedical Engineering*, December 6, 1–11.

Dierssen, Mara. 2012. “Down Syndrome: The Brain in Trisomic Mode.” Nature Reviews. Neuroscience 13 (12): 844–858.

Dunn, Lauren N., Brian F. Niemeyer, Neetha P. Eduthan, et al. 2026. “Altered Hepatic Metabolism in Down Syndrome.” Cell Reports 45 (1): 116835.

Face, Hugging. 2025. “Nvidia/llama-Embed-Nemotron-8b · Hugging Face.” https://huggingface.co/nvidia/llama-embed-nemotron-8b.

Face, Hugging. 2026. “Nvidia/llama-Nemotron-Rerank-1b-v2 · Hugging Face.” https://huggingface.co/nvidia/llama-nemotron-rerank-1b-v2.

Gillenwater, Lucas A., Matthew D. Galbraith, Angela L. Rachubinski, et al. 2024. “Integrated Analysis of Immunometabolic Interactions in Down Syndrome.” Science Advances 10 (50): eadq3073.

“Gpt-oss:120b-Cloud.” 2025. https://ollama.com/library/gpt-oss:120b-cloud.

Huang, Kexin, Serena Zhang, Hanchen Wang, et al. 2025. “Biomni: A General-Purpose Biomedical AI Agent.” In bioRxivorg. June 2. 10.1101/2025.05.30.656746.

“LangChain: Observe, Evaluate, and Deploy Reliable AI Agents.” 2022. https://www.langchain.com/.

Lewis, Patrick, Ethan Perez, Aleksandra Piktus, et al. 2020. “Retrieval-Augmented Generation for Knowledge-Intensive NLP Tasks.” In arXiv [cs.CL]. May 22. arXiv. 10.48550/arXiv.2005.11401.

Lin, C-Y. 2004. “ROUGE: A Package for Automatic Evaluation of Summaries.” In Text Summarisation Branches Out: Proceedings of the ACL-04 Workshop.

Liu, Siru, Allison B. McCoy, and Adam Wright. 2025. “Improving Large Language Model Applications in Biomedicine with Retrieval-Augmented Generation: A Systematic Review, Meta-Analysis, and Clinical Development Guidelines.” Journal of the American Medical Informatics Association 32 (4): 605–615.

Lott, Ira T., and Elizabeth Head. 2019. “Dementia in Down Syndrome: Unique Insights for Alzheimer Disease Research.” Nature Reviews. Neurology 15 (3): 135–147.

Lu, Chris, Cong Lu, Robert Tjarko Lange, et al. 2026. “Towards End-to-End Automation of AI Research.” Nature 651 (8107): 914–919.

McInnes, Leland, John Healy, and James Melville. 2018. “UMAP: Uniform Manifold Approximation and Projection for Dimension Reduction.” In arXiv [stat.ML]. February 9. arXiv. 10.48550/arXiv.1802.03426.

Mégarbané, André, Aimé Ravel, Clotilde Mircher, et al. 2009. “The 50th Anniversary of the Discovery of Trisomy 21: The Past, Present, and Future of Research and Treatment of Down Syndrome.” Genetics in Medicine: Official Journal of the American College of Medical Genetics 11 (9): 611–616.

Mensh, Brett, and Konrad Kording. 2017. “Ten Simple Rules for Structuring Papers.” PLoS Computational Biology 13 (9): e1005619.

Miao, Jiacheng, Joe R. Davis, Yaohui Zhang, Jonathan K. Pritchard, and James Zou. 2025. “Paper2Agent: Reimagining Research Papers as Interactive and Reliable AI Agents.” In arXiv [cs.AI]. October 16. arXiv. 10.48550/arXiv.2509.06917.

“NCBI Entrez Programming Utilities (E-Utilities).” 2009. https://www.ncbi.nlm.nih.gov/pmc/utils/oa/oa.fcgi.

NVIDIA Corporation. 2026. “Nemotron-3-Super.” https://ollama.com/library/nemotron-3-super.

OpenAI. 2026. “Introducing GPT-5.4.” OpenAI. https://openai.com/index/introducing-gpt-5-4/.

Papineni, Kishore, Salim Roukos, Todd Ward, and Wei-Jing Zhu. 2001. “BLEU.” Paper presented the 40th Annual Meeting, 2002/7/7-2002/7/12, Philadelphia, Pennsylvania. Proceedings of the 40th Annual Meeting on Association for Computational Linguistics - ACL ’02. 10.3115/1073083.1073135.

Powers, Rani K., Rachel Culp-Hill, Michael P. Ludwig, et al. 2019. “Trisomy 21 Activates the Kynurenine Pathway via Increased Dosage of Interferon Receptors.” Nature Communications 10 (1): 4766.

Priem, Jason, Heather Piwowar, and Richard Orr. 2022. “OpenAlex: A Fully-Open Index of Scholarly Works, Authors, Venues, Institutions, and Concepts.” In arXiv [cs.DL]. May 4. arXiv. 10.48550/arXiv.2205.01833.

Qu, Yuanhao, Kaixuan Huang, Ming Yin, et al. 2025. “CRISPR-GPT for Agentic Automation of Gene-Editing Experiments.” *Nature Biomedical Engineering*, ahead of print, July 30. 10.1038/s41551-025-01463-z.

Rachubinski, Angela L., Elizabeth Wallace, Emily Gurnee, et al. 2024. “JAK Inhibition Decreases the Autoimmune Burden in Down Syndrome.” eLife 13 (RP99323). 10.7554/elife.99323.3.

Russo, Matthew L., André M. M. Sousa, and Anita Bhattacharyya. 2024. “Consequences of Trisomy 21 for Brain Development in Down Syndrome.” Nature Reviews. Neuroscience 25 (11): 740–755.

Santoro, Jonathan D., Dania Pagarkar, Duong T. Chu, et al. 2021. “Neurologic Complications of Down Syndrome: A Systematic Review.” Journal of Neurology 268 (12): 4495–4509.

Sullivan, Kelly D., Hannah C. Lewis, Amanda A. Hill, et al. 2016. “Trisomy 21 Consistently Activates the Interferon Response.” eLife 5 (e16220). 10.7554/elife.16220.

“The PubMed Central OAI-PMH API.” 2001. https://pmc.ncbi.nlm.nih.gov/tools/oai/.

Topaz, Maxim, Nir Roguin, Pallavi Gupta, Zhihong Zhang, and Laura-Maria Peltonen. 2026. “Fabricated Citations: An Audit across 2·5 Million Biomedical Papers.” Lancet 407 (10541): 1779–1781.

Wang, Zhizheng, Qiao Jin, Chih-Hsuan Wei, et al. 2025. “GeneAgent: Self-Verification Language Agent for Gene-Set Analysis Using Domain Databases.” Nature Methods 22 (8): 1677–1685.

Weijerman, Michel E., and J. Peter de Winter. 2010. “Clinical Practice. The Care of Children with Down Syndrome.” European Journal of Pediatrics 169 (12): 1445–1452.

Zhang, T., V. Kishore, F. Wu, K. Q. Weinberger, and Y. Artzi. 2020. BERTScore: Evaluating Text Generation BERT. International Conference Learning Representations (ICLR 2020).

