## Supplementary material for "An AI-Powered Trisomy 21 Research Assistant": S1_Table_1

| SLNo | CTRL_1 | Original question | Answer | Citation |
|---|---|---|---|---|
| Q1 | Positive | Which clinical conditions are people with Down syndrome at higher risk for? | Individuals with Down syndrome are at high risk for the following conditions: developmental delay, cognitive deficits, congenital heart defects, thyroid dysfunction, autoimmune disorders, respiratory complications, certain leukemias, Alzheimer's disease, and immune dysfunction | |
| Q2 | Positive | Which clinical conditions are people with Down syndrome at lower risk for? | Individuals with Down syndrome are at low risk for many solid tumors and high blood pressure | |
| Q3 | Positive | Why do people with Down syndrome tend to have hyperactive immune systems? | Four interferron receptors, IFNAR1, IFNAR2, IFNGR2, and IL10RB, are located on chromosome 21. These genes are overexpressed in individuals with Down syndrome, leading to increased, chronic interferon signaling, which contributes to increased immune sensitivity and auto-immune risks | |
| Q4 | Positive | Why are people with Down syndrome at high risk of developing Alzheimer's disease? | Amyloid Precursor Protein (APP) is located on chromsome 21 and is upregulated in Down syndrome. The proteolysis of APP generates amyloid beta, a core component of amyloid plaques found in the brain of individuals with Alzheimer's disease | |
| Q5 | Positive | Do individuals with Down syndrome have overactive thyroid glands? | Autoimmune thyroid dysfunction is highly pr | [PMID: 32029743,30298764] |
| Q6 | Positive | Which interferon receptors are triplicated in Down syndrome? | Four of the six interferon receptors are found on human chromosome 21 and thus are triplicated in Down syndrome. These receptor genes are IFNAR1, IFNAR2, IFNGR2, and IL10RB which recognize type I, II and III IFNs respectively. | [PMID: 27472900,10830953,32029743] |
| Q7 | Positive | How is Stat1 signaling affected in Down syndrome? | STAT1 signaling is generally increased due to overexpression of interferon receptors, but the expression can be context dependent. | |
| Q8 | Positive | Which circulating immune cell is consistently depleted in T21 throughout the lifespan? | Individuals with Down syndrome have B cell lymphopenia, which is a condition marked by very low levels of B lymphocytes | [PMID: 40993118] |

| | | | | |
|---|---|---|---|---|
| Q9 | Positive | What are some advantages and caveats of the Ts65Dn and Dp16 mouse models of Down syndrome? | The Dp16 model triplicates the largest number of genes matching human chromosome 21 and is the best option for studying the genetics of Down syndrome; however, the behavioural and congnitive deficits are less pronounced and can be inconsistent. The Ts65Dn model is a robust model for mimicking behaviour and neurological conditions in Down syndrome, but it has triplication of genes that do not match human chromosome 21 and can be challenging to breed. | [PMID: 17412756,2147289] |
| Q10 | Positive | Are individuals with Down syndrome more prone to high blood pressure? | Individuals with Down syndrome typically exhibit lower-than-average blood pressure. | |
| Q11 | Positive | What neurodegeneration is associated with Down syndrome? | Individuals with Down syndrome develop Alzheimer's-like brain pathology by age 40 due to overexpression of the amyloid precursor protein (APP) gene on chromosome 21. Dementia typically manifests with over 50% being affected by age 50. | [PMID: 35676933, 25510383] |
| Q12 | Positive | What type of cancer are individuals with Down Syndrome less prone to developing? | The risk of all major groups of solid tumors, except testicular cancer, are decreased in individuals with Down syndrome. | [PMID: 27031084] |
| Q13 | Negative | How do the PAM50 subtypes in Down syndrome affect treatment options? | No relevant information found in the available literature to answer this question. | |
| Q14 | Negative | What are some drugs that I can use to treat Hypothyroidism in Down syndrome? | No relevant information found in the available literature to answer this question. | |
| Q15 | Negative | How do pine trees relate to the development of Down syndrome? | No relevant information found in the available literature to answer this question. | |
| Q16 | Negative | Do people with Down syndrome have super human strength? | No relevant information found in the available literature to answer this question. | |
| Q17 | Negative | How does the gene FX84A9 drive co-occurring conditions in Down syndrome? | No relevant information found in the available literature to answer this question. | |
| Q18 | Negative | How does chloroplast dysfunction affect co-occuring conditions in Down syndrome? | No relevant information found in the available literature to answer this question. | |

| | | | |
|---|---|---|---|
| Q19 | Negative | Why do all people with Down syndrome have the same diseases and show the same spectrum of co-occuring conditions? | No relevant information found in the available literature to answer this question. |
| Q20 | Negative | Can Ivermectin be used to treat Down syndrome? | No relevant information found in the available literature to answer this question. |
| Q21 | Negative | Are people with Down syndrome more prone to Anthracnose? | No relevant information found in the available literature to answer this question. |
| Q22 | Negative | What behaviorial assarys are used to study Down syndrome in yeast? | No relevant information found in the available literature to answer this question. |
